# Avibactam dose-escalation augments aztreonam-avibactam activity against extensively drug-resistant *Klebsiella pneumoniae* persisters

**DOI:** 10.64898/2026.11.20.767394

**Authors:** Jack F. Klem, Jan Naseer Kaur, Yang Liu, Lu Han, Sarah J. Dinardo, Shivali Kapoor, Patricia N. Holden, Rae Zhu, Jimmy Tangchittsumran, Brian T. Tsuji

## Abstract

**Objectives:** Strategies to address persistence during β-lactam/β-lactamase inhibitor (BL/BLI) therapy remain undefined. To address this clinical gap, we explored increasing AVI exposure throughout aztreonam/avibactam (ATM-AVI) administration to enhance combination activity and suppress extensively drug-resistant (XDR) *Klebsiella pneumoniae* persisters.

**Methods:** 24h time courses were profiled against three XDR *K. pneumoniae* isolates with exposure to increasing concentrations of AVI as monotherapy and in combination with ATM. Sigmoidal E_max_ modeling was conducted to characterize the antibacterial activity of AVI against the tested isolates. In hollow fiber infection model (HFIM) experiments, four ATM-AVI continuous infusion regimens were administered with successively higher AVI dosing against *K. pneumoniae* Nevada over 168h, followed by 120h reversion experiments. Scanning electron microscopy (SEM) provided visualization of morphologic responses to treatment.

**Results:** Time-kill assays, supported by E_max_ modeling, demonstrated AVI concentration-dependent reductions in total bacterial burden over 8h across tested isolates. All ATM-AVI HFIM regimens reduced baseline bacterial burdens below detection over 168h of treatment. However, population reversion to >9.25 log_10_ CFU/mL was observed following treatment with regimens simulating AVI dosed at 1.5 and 6 g/day. Conversely, populations remained suppressed below detection through the 288h endpoint following treatment with proof-of-concept regimens simulating AVI dosed at 24 and 96 g/day. 2h SEM imaging revealed a negative correlation between cell elongation and AVI exposure, which was hypothesized to influence persister suppression and reversion outcomes.

**Conclusion:** This work supports AVI dose escalation to enhance ATM-AVI treatment efficacy and establishes a novel framework by leveraging BLIs to suppress persistence associated with BL/BLI therapy.

## INTRODUCTION

Carbapenem-resistant Enterobacterales (CRE) continue to impose a significant burden on healthcare systems worldwide, with carbapenem-resistant *Klebsiella pneumoniae* occupying the top position in the 2024 WHO Bacterial Priority Pathogens List^1, 2^. Despite growing concerns, resistance is outpacing regulatory approval of new therapies, a situation further exacerbated by the increasing prevalence of metallo-β-lactamase (MBL)-producing pathogens, which are considered notoriously difficult to treat^3, 4^.

With limited therapeutic options available, clinicians have become increasingly reliant on β-lactam/β-lactamase inhibitor (BL/BLI) combinations to treat CRE infections. Most notably, aztreonam/avibactam (ATM-AVI) occupies an important clinical niche as a first-line therapy for treating infections caused by MBL-producing, extensively drug-resistant (XDR) Gram-negatives^5,6^. ATM-AVI is distinct from other BL/BLIs in that, at its core, ATM represents one of few β-lactams inherently stable against MBL-mediated hydrolysis^5–8^. Meanwhile, AVI was the first diazabicyclooctane (DBO) BLI marketed for clinical use, providing broad-spectrum inhibition of serine β-lactamases capable of hydrolyzing ATM^9, 10^. In combination, AVI restores activity to ATM, rendering it effective in the setting of MBL and serine β-lactamase co-expression^8, 11–13^.

However, despite its clinical utility, microbiological persistence has been reported following ATM-AVI treatment in patients with complicated intra-abdominal infections and hospital-acquired/ventilator-associated pneumonia, as indicated in secondary outcomes of the Phase III REVISIT Trial^8^. Although the underlying mechanisms remain undefined, bacterial acquisition of phenotypic adaptations has been proposed as a potential contributor to clinical persistence^14–16^.

As the only clinically available monobactam, ATM exerts its antibacterial activity through high-affinity binding and inactivation of penicillin-binding protein 3 (PBP3), a peptidoglycan transpeptidase involved in septum formation during cellular division^17^. Consequently, this mechanism leads to the formation of filamentous cells, which can fragment following withdrawal of drug pressure and may confound clinical persistence with ATM-based therapies^18–22^.

One approach to addressing persistence with ATM-AVI involves increasing penicillin-binding protein 2 (PBP2) binding to inhibit elongation and complement PBP3 inactivation. Recently, second-generation DBOs have garnered significant attention for their PBP2-driven β-lactam-potentiating properties^9, 10, 23–26^. This enhancer strategy is currently undergoing rigorous clinical development, with several novel BL/BLI candidates nearing or having reached market approval^3, 25, 26^. In the setting of enzyme-mediated resistance, DBOs evade enzymatic inactivation and reversibly inhibit β-lactamases, allowing excess molecules to bind PBP2^9, 23–28^. Due to lower affinity for PBP2 relative to its successors, the antimicrobial activity of AVI was never considered clinically applicable^9, 23, 28^. Given its DBO chemical characteristics, alongside its widespread use and favorable safety profile, exploration of high-dose AVI regimens presents a promising strategy to effectively inhibit PBP2 and combat persistence^28, 29^.

The association of persistence with current dosing guidelines calls for therapeutic optimization of ATM-AVI to improve treatment outcomes and preserve its utility against XDR pathogens. Here, we hypothesized that increased AVI exposure during treatment with ATM-AVI improves bacterial killing and post-treatment suppression by enhancing PBP2 inhibition to disrupt the elongation of filamentous persister cells. In this study, we evaluated an AVI dose-escalation strategy against several MBL-producing, XDR *K. pneumoniae* clinical isolates using *in vitro* pharmacodynamic models and scanning electron microscopy (SEM). Importantly, this study expands the scope of BL/BLI therapy by exploiting intrinsic BLI activity to target persisters, and it provides a promising rationale for future dose-optimization studies.

## MATERIALS AND METHODS

### Materials and susceptibility testing

Cation-adjusted Mueller-Hinton II Broth (MHB) and Mueller-Hinton II agar (MHA) were used in all experiments, both produced by Becton, Dickinson and Company and supplied by Fisher Scientific (cat. nos. 212322 and 225250, respectively). Minimum inhibitory concentrations (MIC) were determined via broth microdilution in accordance with CLSI standard M07^30^. Three New Delhi metallo-β-lactamase (NDM)-producing *K. pneumoniae* clinical isolates were tested, including *K. pneumoniae* Nevada (BioSample <u>SAMN07265011</u>), S200, and S202^18–20, 22, 28, 31^. ATM and AVI were supplied by AK Scientific (cat. nos. J10266 and 0202AP, respectively).

### Time-kill assays

Time-kill assays were performed in triplicate, as previously described^32^. Overnight cultures grown on MHA were suspended in fresh MHB and incubated in a shaking water bath at 37°C for 1-2h to reach exponential-phase growth. Liquid cultures were then washed with normal saline, optical density standardized at 620 nm, and subsequently diluted to a starting inoculum of 6.00 log_10_ colony-forming units (CFU)/mL in MHB containing antibiotics at the desired final concentrations. Targeted final antibiotic concentrations included ATM 66 mg/L, which corresponds to the *f*C_max_ in critically ill humans treated with a 2 g dose, and AVI ranging from 4.95 to 316.8 mg/L^21^. In accordance with validated AVI dosing protocols in combination with ceftazidime, the AVI concentration series was arrayed around a reference of 9.9 mg/L, corresponding to the simulated steady state *f*C_max_ for a 0.5 g dose administered as a 2h prolonged infusion (2hPI) every 8h (Q8H) in HFIM studies^33^. Resulting 20 mL cultures were incubated in a water bath at 37°C with constant shaking, and plate count analysis was performed at 0, 1, 2, 4, 6, 8, and 24h. 50 µL of serially diluted samples were plated on MHA using a WASP automated spiral plater (Don Whitley Scientific Ltd., UK), followed by 24h incubation at 37°C. Colonies were then counted using an @COLyte 3HD plate reader (Synbiosis, UK) to generate CFU/mL trends over 24h. For mean AVI monotherapy data, 0-8h reductions in area under the bacterial killing curve (ΔAUC_0-8_) were calculated using the trapezoidal method and expressed relative to corresponding growth controls.

### E_max_ modeling

For each isolate and AVI concentration, the area under the 0-24h time-kill curve (AUC_0-24_) was calculated from mean viable counts (log_10_ CFU/mL) across triplicate AVI monotherapy assays. Each AUC_0-24_ was normalized to that of the corresponding drug-free growth control to obtain a treated-to-control AUC ratio (growth control = 1). A sigmoidal inhibitory E_max_ model was fitted separately for each isolate to describe the relationship between AVI concentration and the AUC ratio. The estimated parameters were the maximal fractional reduction in AUC_0-24_ relative to the growth control (E_max_), the AVI concentration producing half of this reduction (EC_50_), and the Hill coefficient (γ), which describes the steepness of the concentration-response curve. All analyses were performed in R (version 4.5.3; R Foundation for Statistical Computing, Vienna, Austria), with model parameters estimated by nonlinear least-squares regression using the stats package (version 4.5.1).

### Hollow fiber infection model

In HFIM experiments, cellulosic cartridges (FiberCell Systems, Inc.; C3008) were inoculated with *K. pneumoniae* Nevada at an average of 7.37 log_10_ (SD = 0.11) CFU/mL, as previously described^18, 34, 35^. The models were treated with respective antibiotic regimens over 168h, followed by a 120h reversion phase to confirm or rule out sustained persister suppression beyond our traditional 48h window^18–22^. Persistence was assessed by determining MICs for reverted samples collected at 216h. Preserved susceptibilities to ATM-AVI were indicative of persistence rather than resistance in reverted populations^15^. The flow rate was maintained at 0.8 mL/min throughout the experiment to simulate 2h half-lives for ATM and AVI based on previously validated run conditions^7, 33, 36^. Continuous infusion (CI) administration was selected to maximize time above the MIC and to prevent troughs, which we postulated could promote the establishment of persister subpopulations^37^. The following regimens were simulated in the set-up design:

i. Growth Control^18, 19, 22^
ii. ATM 6 g/day
iii. AVI 96 g/day
iv. ATM-AVI 6/1.5 g/day
v. ATM-AVI 6/6 g/day
vi. ATM-AVI 6/24 g/day
vii. ATM-AVI 6/96 g/day

Regimen IV served as the reference standard and was AUC-adjusted for CI to achieve equivalent ATM and AVI exposures, based on previously validated dosing recommendations for ATM in combination with ceftazidime/AVI^33^. To investigate the effect of increasing AVI exposure, the ATM dose was fixed at 6 g/day, while the AVI dose was sequentially increased fourfold from 1.5 to 96 g/day (regimens IV – VII). Regimens VI and VII represent proof-of-concept dosing targets used to gain mechanistic insights into the effects of high-dose AVI. Pharmacodynamics were assessed through quantitative bacterial plate counts at indicated timepoints from 0 to 288h.

### Scanning electron microscopy

Morphological responses to treatment were visualized through scanning electron microscopy (SEM) of samples collected from HFIM cartridges at indicated timepoints, as previously described^21^. Cells were fixed in 2.4% glutaraldehyde (Electron Microscopy Sciences, USA) and refrigerated for at least 12h before processing. Preserved samples were syringe-filtered onto 0.2 µm polycarbonate Nuclepore membranes (Whatman, USA) and dehydrated through graded ethanol washes (30%, 50%, 70%, 80%, 90%, 95%, 97.5%, 2x 100% v/v). Following dehydration, the membranes were incubated in hexamethyldisilazane (HMDS) for 5 minutes and transferred to empty Petri dishes to dry completely. Samples were then carbon-coated using a Denton 502 evaporator and imaged using a Hitachi SU70 FE-SEM at an accelerating voltage of 2.0 kV (Hitachi High-Tech America, Inc., USA).

## RESULTS

### Baseline Susceptibility Screening

Baseline susceptibility profiles of the three targeted *K. pneumoniae* isolates were determined through MIC testing (**Table 1**). All isolates exhibited high-level resistance to ATM and AVI monotherapies but were susceptible to ATM-AVI combination therapy. Based on these results and known resistance profiles, all three isolates were characterized as XDR and included in subsequent experimentation.

**Table 1.**
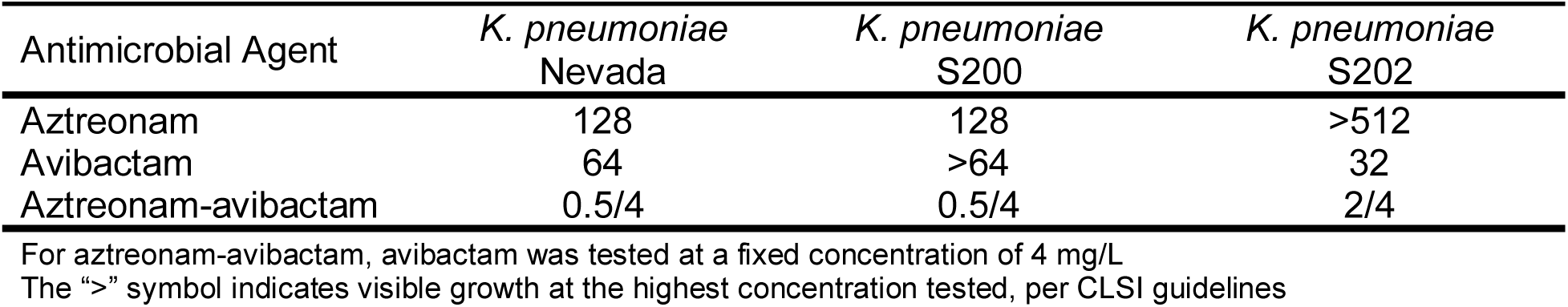
Relevant minimum inhibitory concentrations (MIC, mg/L) against evaluated isolates.

### Time-kill assays and E_max_ modeling

In time-kill assays, AVI monotherapy failed to reduce baseline bacterial burdens by >1.0 log_10_ CFU/mL over 24h for all isolates tested. However, despite subsequent regrowth, concentration-dependent reductions in total bacterial burden were observed over the first 8h of treatment relative to corresponding growth controls (**Figure 1**). Specifically, ΔAUC_0-8_ generally increased across AVI concentrations of 4.95–316.8 mg/L, ranging from 3.3 to 24.5, 0.7 to 26.2, and 0.7 to 26.7 log_10_ CFU·h/mL for *K. pneumoniae* Nevada, S200, and S202, respectively. Regrowth by 24h suggested that all three of the isolates either contained subpopulations that were highly resistant to AVI monotherapy or experienced adaptive resistance over the 24h duration of exposure. Sigmoidal E_max_ modeling results were derived from mean triplicate AVI monotherapy data over 24h (**Figure 1**). AUC_0-24_ ratios were plotted against AVI concentration and yielded EC_50_ estimates of 8.19, 764.52, and 15.99 mg/L for *K. pneumoniae* Nevada, S200, and S202, respectively. AVI exhibited the greatest potency against *K. pneumoniae* Nevada and the lowest potency against *K. pneumoniae* S200, which was supported by its lowest and highest EC_50_ values, respectively. The significantly larger E_max_ threshold observed against *K. pneumoniae* S200 relative to the other isolates tested suggests that this strain may exhibit the greatest cumulative expression of β-lactamases, requiring markedly higher AVI exposure levels to outcompete enzymes and saturate PBP2 binding sites. Altogether, all isolates demonstrated concentration-dependent responses to AVI, albeit to varying degrees, providing further evidence of AVI’s intrinsic antimicrobial activity.

**Figure 1.**
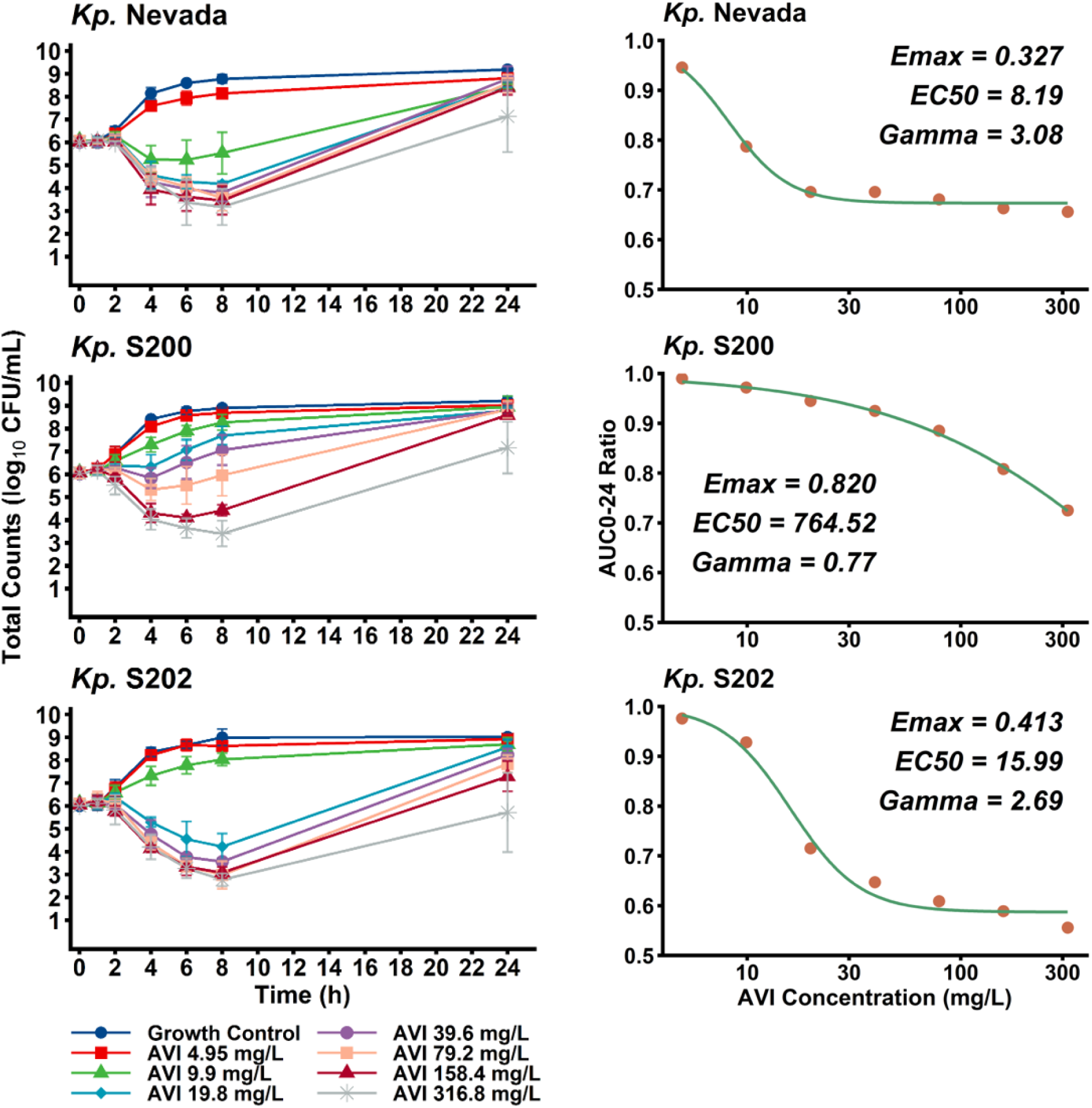
24h time courses against *K. pneumoniae* Nevada, S200, and S202 (**left**) with corresponding sigmoidal E_max_ modeling curves (**right**). All bacterial cultures were diluted to an initial inoculum of ∼6.00 log_10_ CFU/mL and exposed to AVI ranging from 4.95 to 316.8 mg/L. Sigmoidal E_max_ modeling provided quantification of strain-specific responses to escalating AVI exposure. Experiments were performed in triplicate, and standard deviations are indicated by error bars.

In ATM-AVI combination arms, clinically achievable AVI exposure was sufficient to produce >2.00 log_10_ CFU/mL reductions in baseline bacterial burdens. However, increasing AVI concentrations were generally associated with lower total counts at 24h (**Figure 2**). *K. pneumoniae* S200 and S202 exhibited greater susceptibility to combinations compared to *K. pneumoniae* Nevada, which saw the most pronounced count separation at 24h, ranging from 3.7 log_10_ CFU/mL to below the limit of detection. Interestingly, *K. pneumoniae* Nevada was the most difficult to treat with ATM-AVI combinations despite being the most sensitive to AVI monotherapy. This concentration-dependent response extended beyond the concentration associated with AVI’s near-maximal effect (∼30 mg/L, sigmoidal plateau), which signaled a mechanism that may extend beyond baseline enzyme outcompetition and binding site saturation. We inferred that this secondary mechanism could be attributed to suppression of ATM-mediated persistence with elevated AVI exposure, which prompted us to explore this isolate in the HFIM.

**Figure 2.**
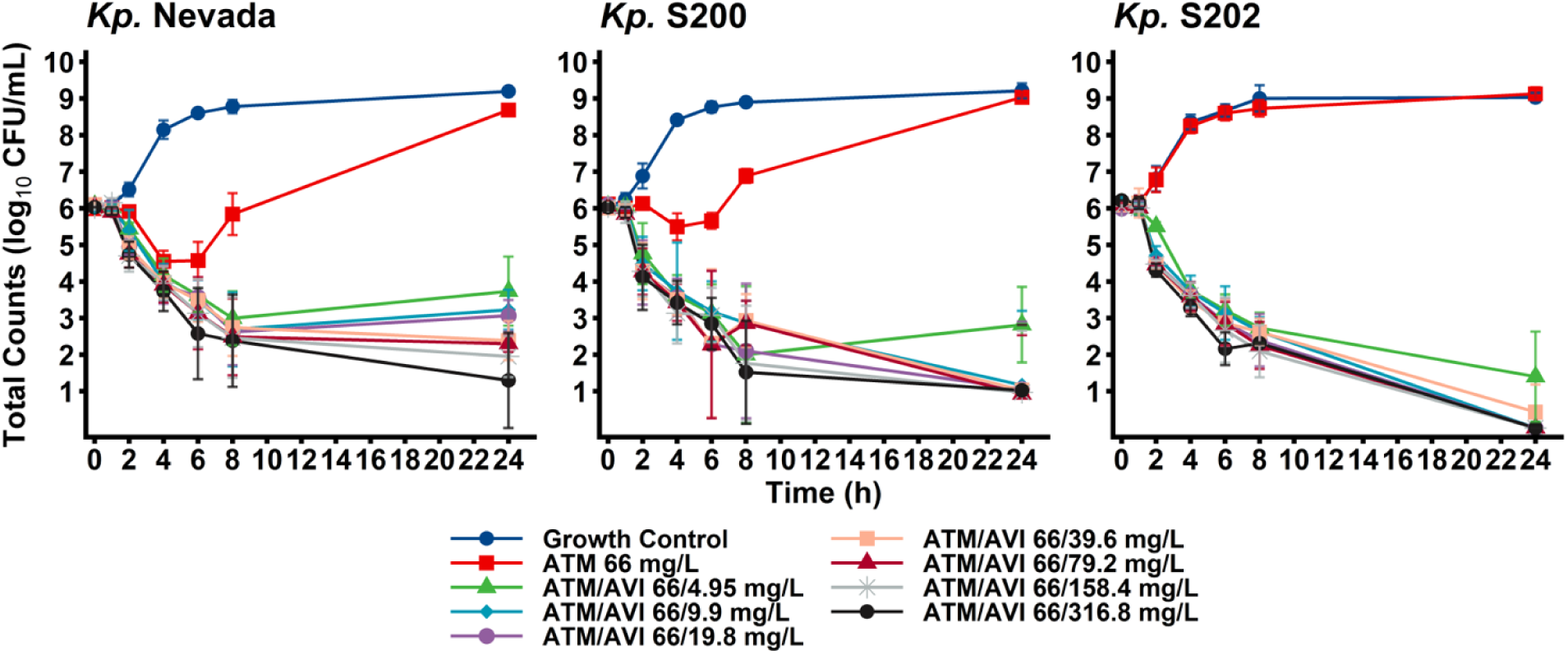
Evaluating AVI in the presence of ATM (66 mg/L, fixed) against *K. pneumoniae* Nevada, S200, and S202 in time-kill assays. Results demonstrated ATM-AVI synergy over 24h with a notable concentration-dependent response observed in 24h counts for *K. pneumoniae* Nevada. Experiments were performed in triplicate, and standard deviations are indicated by error bars.

### Hollow fiber infection model

In the HFIM, ATM monotherapy (regimen II) performed as expected, with total counts reaching the system’s carrying capacity within 24h, consistent with previous results ^18, 19, 22^ (**Figure 3**). AVI monotherapy (regimen III) produced an initial 3.20 log_10_ CFU/mL total count reduction within 6h, followed by regrowth to 9.25 log_10_ CFU/mL by 48h. All ATM-AVI regimens (regimens IV – VII) reduced total counts below the limit of detection by 72h, with sustained population suppression through 168h (reductions from baseline of 7.41, 7.35, 7.20, and 7.26 log_10_ CFU/mL, respectively). Treatment with ATM-AVI 1/1x and 1/4x (regimens IV and V) resulted in population regrowth following drug discontinuation, with counts exceeding 9.25 log_10_ CFU/mL by 216h. Persistence was confirmed through 216h MIC assays, which highlighted the preservation of original susceptibilities to ATM-AVI in the reverted populations. Notably, ATM-AVI 1/16x and 1/64x (regimens VI and VII) maintained suppression below detection through 216h, 48h after drug discontinuation. Consistent with our hypothesis, combination treatment involving higher doses of AVI was associated with suppression of persister subpopulations following treatment cessation.

**Figure 3.**
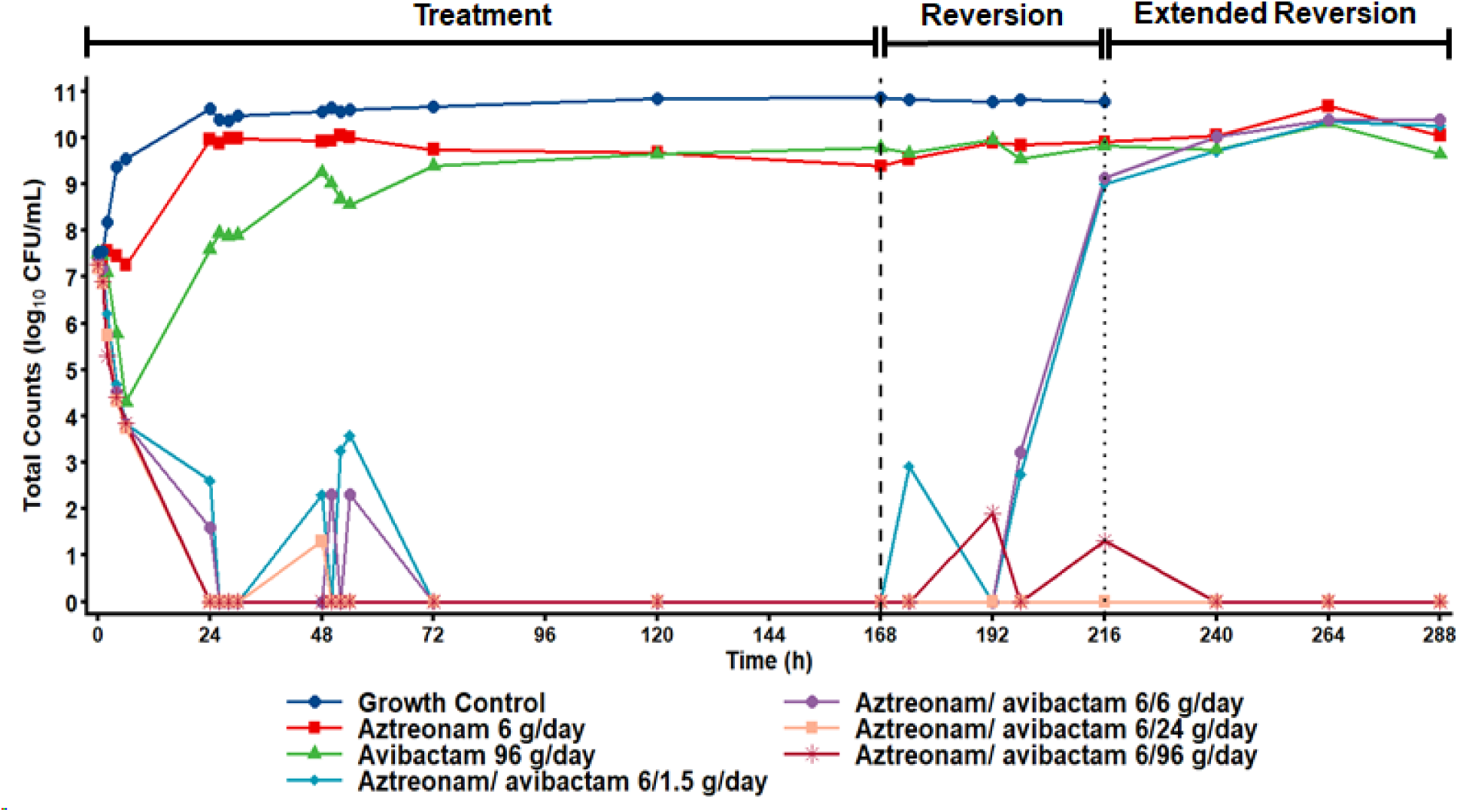
Tracking viable plate counts for *K. pneumoniae* Nevada with an initial inoculum of ∼7.3 log_10_ CFU/mL in a 288h HFIM during and after treatment with indicated ATM-AVI CI regimens. Antimicrobial administration was discontinued after 168h (**long-dashed line**), and the experiment was continued beyond our standard 48h reversion period (**short-dashed line**) to a 288h endpoint, with viable count sampling performed every 24h to assess population viability over an extended five-day duration of favorable growth conditions.

To substantiate this finding, we extended the reversion period by three days (**Figure 3**). Accordingly, both populations treated with ATM-AVI 1/16x and 1/64x (regimens VI and VII) failed to regrow to carrying capacity prior to the 12-day endpoint, further highlighting the persister-suppressing effects of high-dose AVI.

### Scanning electron microscopy

As expected, ATM and AVI monotherapy produced filamentous and spheroplast phenotypes, respectively, throughout the HFIM treatment window. Conversely, treatment with ATM-AVI combination therapy (regimens IV – VII) produced intermediate bulging, elongated, rod-shaped cells within 6h of treatment. A predominance of cellular debris was observed in all combination samples by 168h, which was consistent with undetectable counts (**Figure 4**). Interestingly, treatment with 1/1x and 1/4x resulted in persister proliferation, supported by 216h imaging of mixed rod-filament populations, whereas samples treated with 1/16x and 1/64x showed sustained predominance of cellular debris through 216h. Most notably, 2h imaging revealed more apparent cell elongation in samples treated with 1/1x and 1/4x (regimens IV and V) compared to those treated with 1/16x and 1/64x (regimens VI and VII) (**Figure 5**). Here, increasing AVI exposure resulted in notable attenuation of early filamentation, which was associated with persister suppression following treatment cessation. These findings suggest that inhibition of cellular elongation could compromise the establishment of persister subpopulations during ATM-AVI therapy.

**Figure 4.**
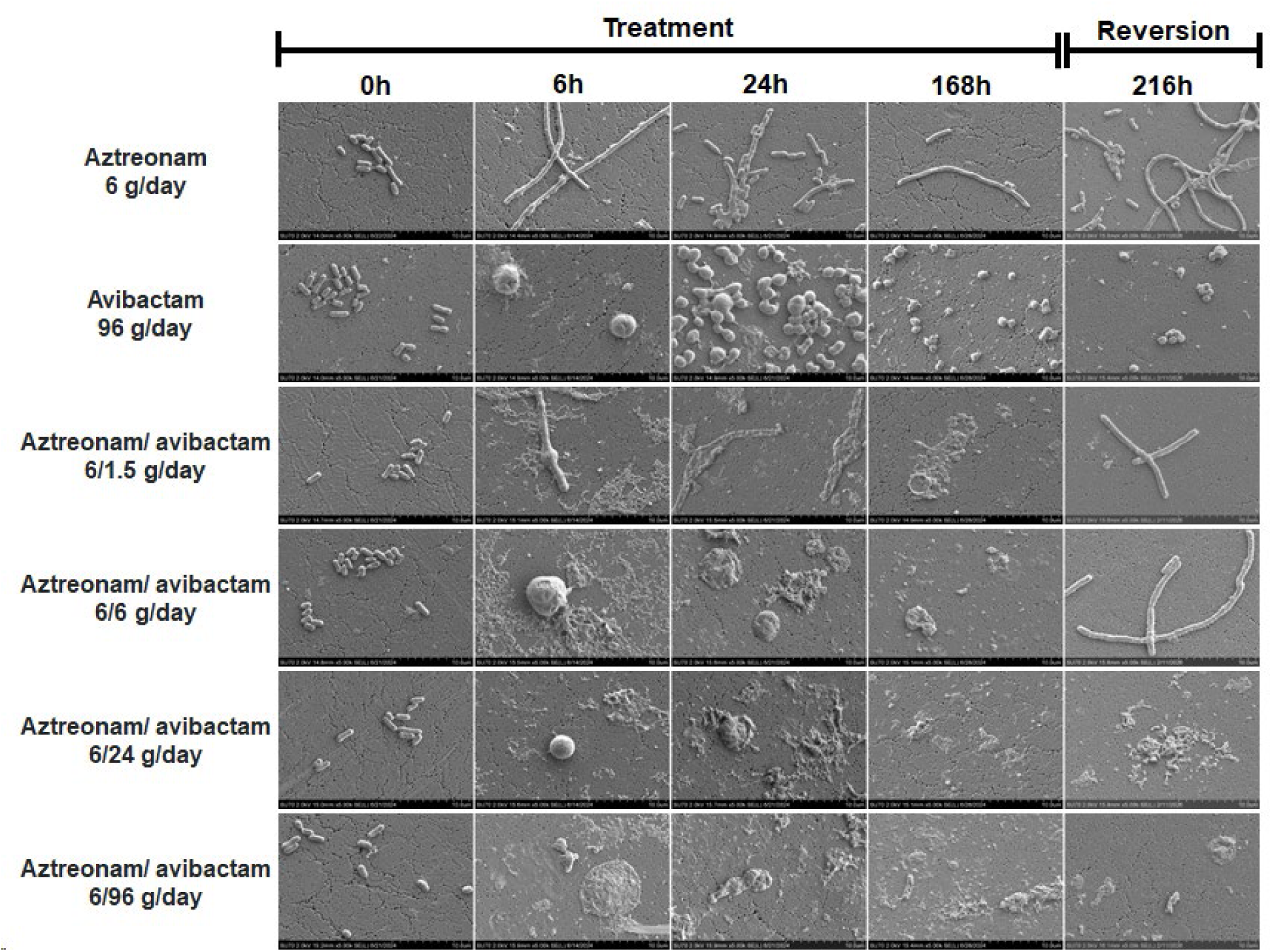
Representative SEM imaging of *K. pneumoniae* Nevada depicting morphological responses during and after treatment with indicated HFIM regimens. Images were captured at 5000x magnification using a Hitachi SU70 FE-SEM with an acceleration voltage of 2.0kV.

**Figure 5.**
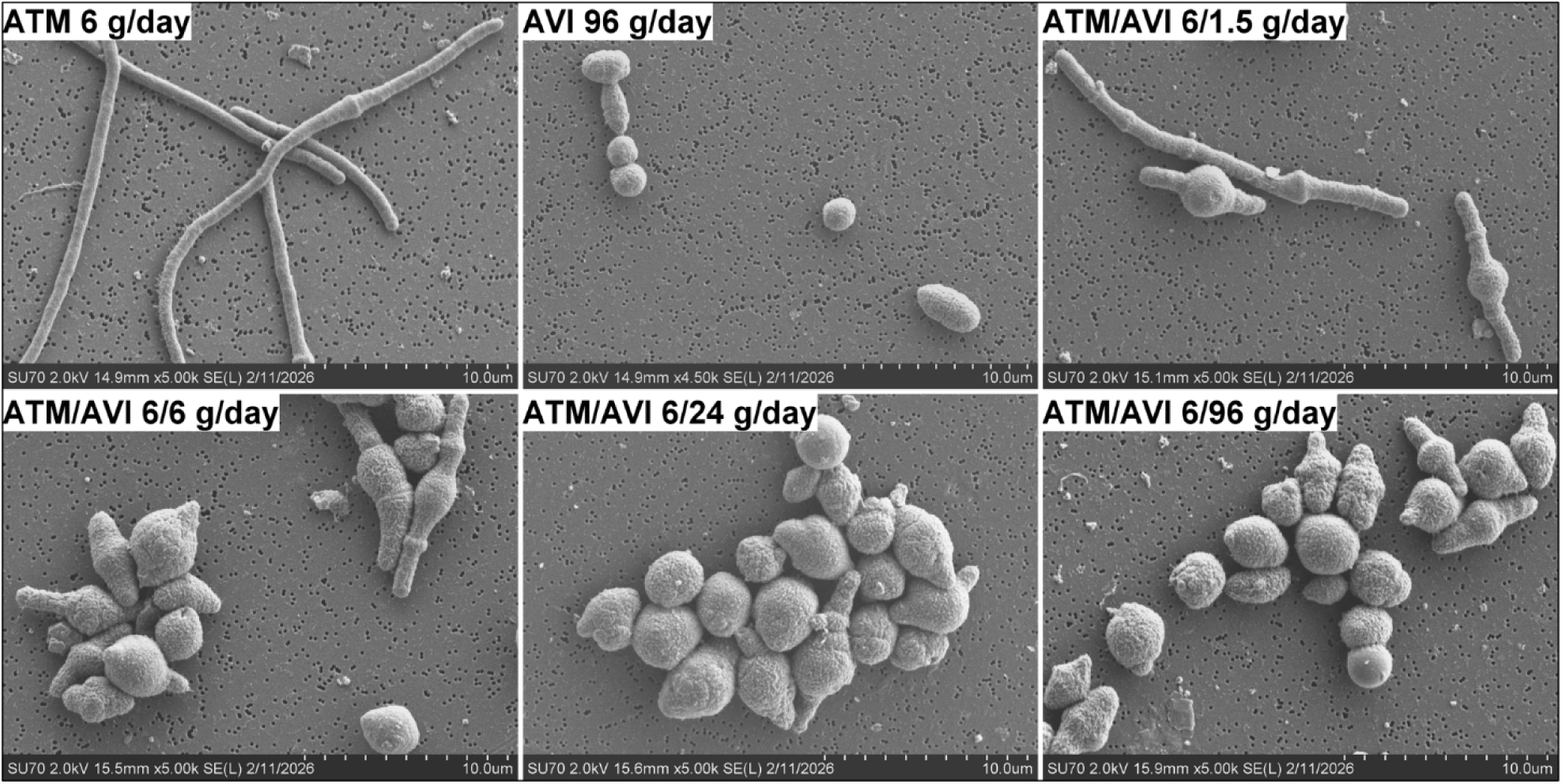
SEM images showing a negative correlation between cellular elongation and AVI exposure. *K. pneumoniae* Nevada samples were extracted from the HFIM at the 2h time point and imaged at 5000x magnification using a Hitachi SU70 FE-SEM with an acceleration voltage of 2.0kV.

## DISCUSSION

Clinicians lack guidance on how to address persistence during and following BL/BLI therapy. Given the low-density nature of persisters, developing therapeutic solutions targeting these subpopulations has historically been difficult^14, 15, 38, 39^. Here, we demonstrated that escalating AVI exposure during ATM-AVI therapy disrupts cell elongation and suppresses persistence in XDR *K. pneumoniae,* consistent with enhanced PBP2 inhibition.

AVI has previously demonstrated *in vitro* activity against MBL-producing Gram-negatives, including *K. pneumoniae* Nevada, at concentrations exceeding those achieved clinically^28^. The current study expands upon these findings by evaluating the effects of increasing AVI exposure alone and in combination with ATM against additional XDR clinical isolates. Despite strain-specific differences in response magnitude, increasing AVI concentrations consistently enhanced antibacterial activity across all tested isolates over 8 h, as supported by E_max_ modeling. Furthermore, significant AVI synergy with ATM was observed across isolates, suggesting that AVI-mediated enhancement of ATM activity may involve both increased β-lactamase inhibition and potential PBP2 engagement.

In HFIM experiments, elevated AVI dosing was associated with suppressed persister subpopulations, supported by undetectable bacterial counts during the extended reversion phase and SEM imaging of cellular debris following treatment discontinuation. Interestingly, higher AVI dosing was also associated with markedly reduced cellular elongation at the beginning of therapy relative to standard dosing, as observed in 2h SEM imaging. This finding may be attributed to enhanced PBP2 inhibition with heightened AVI exposure, and it is of particular interest given ATM’s propensity to induce filamentous persister cells^18–22, 28, 40^.

Inhibition of cell elongation has been documented with combined exposure to PBP2 and PBP3-binding β-lactams^19, 41^. Recently, this morphologic response has also been characterized in MBL-producing *Escherichia coli* isolates exposed to cefepime/zidebactam, demonstrating how PBP2-binding BLIs can lead to the formation of intermediate, bulging, elongated cells when administered alongside PBP3-binding cephalosporins^42^. Over time, these intermediate cells continue to swell until they are no longer able to compensate for the growing osmotic pressure, resulting in lysis via a PBP2 inhibition-dependent mechanism, as previously described^43–45^. This “osmotic overwhelming” mechanism holds therapeutic promise because unlike filamentation, which allows for DNA replication and cellular expansion, it provides a direct pathway to irreversible cell death and represents a potential strategy to overcome PBP3-mediated persistence^21, 46^.

It should be noted that persistence has been reported during monotherapy with PBP2-binding agents^47, 48^. Like our previous work demonstrating tolerance and reversion of filamentous persisters, Cai et al. and Cross et al. observed a similar phenomenon in drug-treated cells exhibiting a spheroplast phenotype characteristic of PBP2 inactivation^21, 47, 48^. Notably, these studies involved exposure times of ≤24h. We hypothesize that dynamic, humanized, prolonged ATM-AVI exposure (i.e., 168h) is sufficient to lyse drug-tolerant spheroplasts. Nevertheless, persistence associated with PBP2-inhibited cells, as well as the threshold at which spheroplasts become non-viable, warrants further study. We propose AVI dose escalation as a strategy to exceed this threshold to lyse spheroplasts that may otherwise contribute to persistence.

To date, elevated AVI dosing beyond what is required for enzymatic inhibition has not been thoroughly investigated. Current ATM-AVI dosing recommendations call for a maintenance AVI dose of 2 g/day; however, there is clinical evidence that the dose may be safely increased by at least 1.5-fold^7, 29^. Preliminary data from our group also suggest that a 6 g/day target may be clinically achievable based on murine extrapolation studies^49^. Although 6 g/day failed to suppress persisters in the HFIM, less cell elongation was observed at 2h relative to the 1.5 g/day treatment arm. Given the *in vitro* nature of the HFIM, we hypothesize that AVI dosed at 6 g/day is sufficient to suppress persisters enough for the immune system to eradicate remaining viable cells. Moreover, Ulloa et al. demonstrated that AVI sensitizes *K. pneumoniae* to clearance by the innate immune system, providing a pathway for additional mechanistic synergy to overcome persistence^24^.

We acknowledge several limitations of our study. As previously mentioned, the *in vitro* nature of the current data warrants further consideration of AVI dose escalation in *in vivo* models to fully elucidate its place in BL/BLI therapy. Additional humanized exposure-toxicity studies are needed to establish clinically achievable thresholds to inform AVI dose escalation. Genomic profiling of tested isolates and reverted populations could provide insights into molecular mechanisms associated with AVI-related persister suppression and identify additional pathways for therapeutic intervention. BLI dose escalation may prove to be effective in enhancing additional BL/BLI combinations, meriting comprehensive study across current therapies. Crucially, expansion of this strategy to more Gram-negative pathogens should be considered to cover the broad spectrum of CRE urgent threats.

Overall, this study expands upon our previous work by leveraging AVI’s intrinsic antimicrobial properties to target filamentous persisters. From a clinical perspective, it challenges the current paradigm for BL/BLI therapy and establishes a foundation for future PK/PD exploration of BLIs as persister-selective therapeutics.

## Acknowledgements

We thank Peter Bush and the University at Buffalo South Campus Instrumentation Center for training and assistance with Scanning Electron Microscope image acquisition.

## Transparency and declarations

The authors of this manuscript have no conflicts of interest to disclose.

